# 3D Co-culture Model on the Role of Vimentin in Notch Signaling and Vascularization

**DOI:** 10.1101/2023.03.02.530837

**Authors:** Khalil Dayekh, Kibret Mequanint

## Abstract

The Notch signaling pathway is a conserved pathway that is central in vascular tissue development and pathology. Because this pathway controls such important events, it is regulated at multiple steps of its cascade, such as post-translational modification of its ligand and receptor. Recent studies have suggested regulation of the Notch signaling by a pulling force to be required to activate Notch signaling. In this exploratory study, 3D fibrin gels were used as a co-culture system of endothelial cells and 10T1/2 cells to assess whether vimentin is implicated in the regulation of Notch signaling and neovascularization. The results show that 10T1/2 cells increase the expression of Hes-1, Hes-5, and Acta2 during co-culture with human coronary artery endothelial cells (HCAECs) and that vimentin knock-down using siRNA partially reduced the expression under static conditions. On the other hand, while the same trend was observed for Hes-5 under dynamic conditions, Acta2 was overexpressed, and vimentin knock-down did not affect its expression levels. Moreover, the development of newly formed micro-vessels is observed in 3D fibrin gels in the presence of VEGF but could not be formed when vimentin expression was knocked down. These results suggest that vimentin plays a secondary role in Notch signaling; however, it is essential for neovascularization.

## Introduction

Vimentin is a type of intermediate filament protein and is a major component of the cytoskeleton. It is widely expressed in many tissues, including the brain, lungs, liver, gastrointestinal tract, kidneys etc. The vimentin monomer is a 466 amino acid protein and has a molecular weight of about 53 kDa (NCBI Reference Sequence: NP_003371.2). During assembly, two vimentin polypeptides align and bind, forming a dimer. This step is followed by the lateral binding of two dimers to form a tetramer, then eight tetramers bind side-by-side, forming the unit-length fiber (ULF). The ULFs are the basic building blocks of the vimentin intermediate fibers that join end to end^1^ (Figure 1.). The assembly of vimentin fibers is regulated by the phosphorylation of serine residues, which has been shown to disassemble the fibrillar structure of vimentin^2^. Vimentin mainly functions as a structural support protein, and earlier studies in vimentin knock-out mice showed that its absence has no effect on the survival of mice that showed no obvious abnormalities^3^. On the other hand, more recent studies have shown that it is involved in important cellular processes. As an example, vimentin plays a role in cell adhesion due to its ability to interact with integrin at focal adhesion sites^4^. Furthermore, vimentin knock-down in alveolar epithelial cells using shRNA has been shown to reduce their mobility and consequently affect wound-closure rates in an in vitro model of lung injury, and ectopic expression of vimentin reversed those effects^5^. Vimentin has also been implicated in other important processes, such as proliferation^6^ and differentiation^7^.

**Figure 1.**
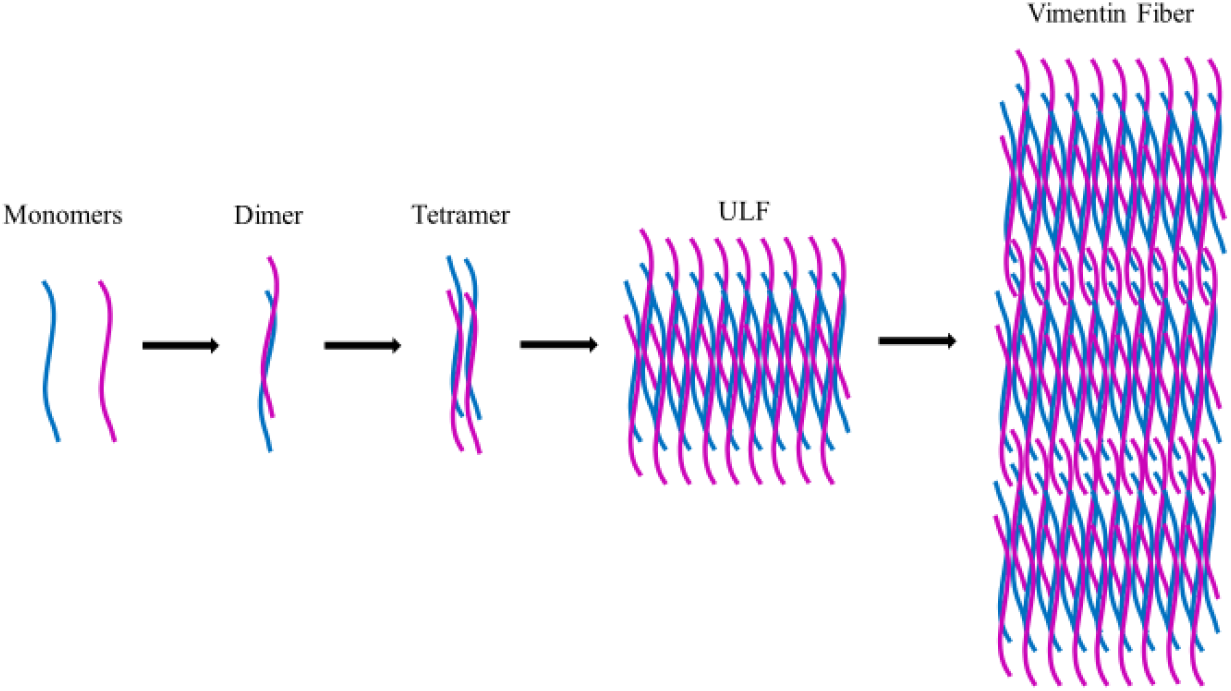
Schematic representation of the assembly of vimentin intermediate fibers.

Interestingly, vimentin has been shown to play a role in regulating signaling pathways, for example, by transporting kinases from one part of a cell to another^8^, or localizing receptors to the cell surface^9^. Of importance to this research, new evidence suggests that the Notch signaling pathway requires a force pulling on the ligand-bound receptor to activate the signal^10, 11^. Naturally, the pulling force has to be exerted by the cytoskeleton component, and while some studies suggest that the actin fibers are involved^11^, other studies have pointed to vimentin^12, 13^. Although the Notch signaling pathway is known to be a prominent regulator of vascular development and homeostasis^14-16^, it is still not a well-understood pathway due to its complex regulation and context dependence. In this exploratory c0-culture study, the role of vimentin in regulating the Notch pathway signaling and its role in neovascularization is inspected.

## Materials and Methods

### Cell culture and tissue fabrication

Embryonic multipotent mesenchymal progenitor cells (10T1/2 cells) (ATCC) were maintained in Dulbecco’s Modified Eagle’s Medium (DMEM) (Gibco) supplemented with 5% v/v Fetal Calf Serum (Gibco) and 1% v/v penicillin/streptomycin (Life Technologies). Human coronary artery endothelial cells (HCAEC) (Lonza) were maintained in EGM^™^ -2 Endothelial Cell Growth Medium-2 BulletKit^™^ (Lonza). Spent media was exchanged with fresh media every three days for 10T1/2 cells and two days for HCAECs. Cells were passaged when confluency reached around 80%. For certain experiments, HCAECs were treated with 3 ng/mL VEGF, EGF, or bFGF or transfected with vimentin siRNA for 3 days prior to co-culture with 10T1/2 cells. Some experiments were performed under dynamic conditions indicating that the culture plates were incubated on an orbital shaker for 30 minutes at a speed of 120 rpm for a period of 3 days.

To fabricate engineered vascular tissues, cultured cells were trypsinized with trypsin-EDTA 0.05% (Thermofisher) for 2 min and then suspended in DMEM. The cells were counted, and the appropriate volume was taken from the cell suspension to give a final cell count of 10 million cells/mL of tissue construct. The cells were then centrifuged at 1200 rpm for 5 min, the supernatant aspirated and cells resuspended in 150 µL of media. To that, 3 µL of 2 M CaCl_2_ and 5 µL of 10 mg/mL ε-aminocaproic acid (ε-ACA) (Sigma-Aldrich), and 1 µL of 1U/µL Thrombin (MP Biomedicals) were added. The cell suspension was kept on ice until it was mixed with ice-cold 150 µL solutions of 6 mg/mL bovine fibrinogen (MP Biomedicals) to give a final concentration of 3 mg/mL fibrinogen per construct. Right after mixing the two solutions, the mixture was transferred into a 5 mL round-bottom tube. The tubes were then transferred to an incubator at 37 °C for 1.5 h for crosslinking. A 3 mL volume of prewarmed DMEM was then added to the tube. The next day, HCAECs were trypsinized using the same procedure, and 1.0 × 10^4^ cells were added to the cultured tissues to form an endothelial cell layer, followed by overnight incubation of the tissues to allow the endothelial cells to adhere. After that, the tissues were taken out of the tubes and cultured in a 50:50 mixture of DMEM: EGM in a culture plate.

### Capillary formation assay in fibrin gel

Tissue plugs made of 3 mg/mL fibrinogen and containing a mixture of 10T1/2 (10 million cells/mL) and HCAECs or siRNA transfected HCAECs (3 million cells/mL) were made in a similar fashion to the tissue preparation protocol above. These plugs were then embedded in 1 mg/mL fibrinogen gels containing 3 ng/mL VEGF or a mixture of 3 ng/mL EGF and bFGF and incubated for a period of 10 days in the presence or absence of 10 µM FOXC2-inhibiting Vimentin effector 1 (FiVe1). The HCAECs cells were stained with cell tracker red (Life Technologies).

### Vimentin knock-down with siRNA

To knock down the expression of vimentin in HCAECs, siRNA reverse transfection protocol was used as described by the manufacturer (Life Technologies) in 24 well plates. The following protocol is based on 1 well of a 24 well plate. In brief, the siRNA-lipofectamine complex was prepared by diluting 6 pmol siRNA in 100 µL of serum and antibiotic-free DMEM. To that, 2 µL lipofectamine RNAiMAX (Life Technologies) was added and mixed gently and then transferred to a plate well. The mixture was incubated at room temperature for 30 minutes. A cell suspension containing 5 × 10^4^ cells /mL was prepared, and 0.5 mL of this suspension was added to each well containing the siRNA-Lipofectamine complex. The plate was incubated for 3 days at 37°C in a CO_2_ incubator before using the cells to allow sufficient time for knock-down. Scrambled siRNA and non-transfected cells were used as control.

### Immunofluorescence microscopy

For 2D studies, cells were seeded in 6-well plates containing a coverslip at a density of 2.5 × 10^5^ cells per well. After 24 h, cells were either left untreated or were treated with control, 3 ng/mL bFGF, EGF, VEGF, or treated with 5 µM of FiVe1 for 3 days. After the culture period, cells were washed with PBS and fixed with 4% paraformaldehyde for 15 min at room temperature (RT). Engineered tissues were fixed overnight in 5 mL tubes, washed with PBS 3 times, and incubated first in a 15% and then in a 30% solution of sucrose at RT until the tissues sunk to the bottom of the tube. The fixed tissues were dabbed using a paper towel to remove the excess liquid. After that, the tissues were immersed with OCT compound (Fisher) and transferred into −80 °C isopropanol. Tissue sections (30 µm thickness) were obtained by using a Leica cryostat (Leica) and placed on microscope slides. The slides were washed with PBS 3 times for 5 min each to remove the OCT compound. Cells/tissue sections were permeabilized with a 0.2% (v/v) Triton x-100 in PBS for 15 min at room temperature and then blocked with 5% BSA in PBS-T for 1 h at RT. The blocking solution was aspirated, and 100 µL of the appropriate primary antibodies anti-Acta2 and Hes-5 antibodies from mouse (Santa Cruz Biotech)) (1:100) in 5% BSA PBS-T were placed on the coverslip and covered with a piece of parafilm, and placed in a humid environment at 4° C overnight. The cells/tissues were washed 2 × with PBS-T and once with PBS for 5 min each, then incubated with the corresponding secondary antibody (Alexa-488 conjugated goat anti-mouse and Alexa-594 conjugated goat anti-rabbit (Life Technologies)) (1:150) in 5% BSA PBS-T for 1 h at RT in the dark. The coverslips were then washed 2 × with PBS-T and once with PBS and incubated with 2 µg/mL DAPI for 5 min, washed with PBS 3 times and mounted with anti-fade mounting media. The images were taken by Zeiss Z1 fluorescent microscope.

### Endothelial cell separation with PECAM beads

For experiments where HCAECs were co-cultured with 10T1/2 cells, the two cell types were separated using PCAM-conjugated magnetic beads^17, 18^. Briefly, the co-cultured cells were washed with HBSS and trypsinized as above. After trypsinization, the cells were suspended in DMEM containing 5% FBS then centrifuged at 1200 rpm at room temperature for 5 minutes. The supernatant was discarded, and the pellet was resuspended in PBS containing 0.1% BSA. PECAM conjugated magnetic beads were used to separate the two cell types by using 100:1 bead to HCAECs ratio. After adding the beads to the cell suspension, the tubes were tumbled end-over-end for 30 minutes at 4 °C. The beads-bound cells were then captured using a magnetic rack and the rest of the suspension containing the 10T1/2 cells were collected and centrifuged at 1200 rpm at 4 °C for 5 minutes. The 10T1/2 cells pellet was collected and washed with ice-cold phosphate-buffered saline (PBS) until further use.

### RNA isolation and qPCR

Pelleted 10T1/2 cells that were separated from HCAECs were collected by centrifugation and 500 µL of Trizol (Life Technologies), and cells were lysed by repeated pipetting. Cells were lysed for 10 min at RT, and chloroform was added at a ratio of 1:5 (chloroform:Trizol), and the samples were vortexed for 15 sec then incubated at RT for 15 min. Samples were then centrifuged at 4 °C and 12000×g for 15 min. The organic phase was discarded, and the aqueous phase was transferred to another Eppendorf tube. Isopropanol was added at a ratio of 1:2 (isopropanol:Trizol) and incubated at RT for 10 min followed by centrifugation at 12000×g for another 10 min at 4 °C. The isopropanol was then aspirated, and the pellet was resuspended in 75% EtOH at a ratio of 1:2 (EtOH:Trizol) and centrifuged at 7500×g for 5 min at 4° C. This last step was repeated twice to wash excess salts. The pellet was air-dried after the EtOH was removed, dissolved in 25 µL of DEPC water, and quantified with nanodrop (Thermo Scientific). 1 µg of total RNA was used to synthesize cDNA using M-MLV reverse transcriptase kit (Promega) using the supplier’s protocol. For qPCR reactions, 1 µL of the formed cDNA was used in 10 µL reactions using the SsoAdvanced universal SYBR green supermix (Bio-rad) according to the manufacturer’s protocol. The qPCR reactions were carried out in a CFX96 Real-Time thermal cycler (BioRad), and GAPDH was used as a reference gene.

### Western blotting

The PECAM separated 10T1/2 cells were harvested using centrifugation and lysed in ice-cold RIPA buffer (50 mM Tris-Cl pH 7.5, 150 mM NaCl, 1 mM EDTA, 1% (v/v) Triton X-100, 0.25% (w/v) sodium deoxycholate and 0.1% (w/v) SDS pH: 7.5) containing protease inhibitor cocktail (Roche) and 1 µM Phenylmethanesulfonyl fluoride (PMSF). The cells were kept on ice for 15 min to allow for lysis to complete. Lysates were centrifuged at 12000 rpm for 15 min. The pellets were discarded, and the supernatants’ protein contents were quantified using the Pierce BCA protein assay protocol (Pierce). Protein samples were resolved by SDS-PAGE and transferred onto a nitrocellulose membrane (Pall life sciences). Blocking the membrane was performed with 5% BSA (Sigma-Aldrich) in PBS with 0.1% Tween-20 (PBS-T) and Western blotted with the Acta2 (1:1000), Hes5 (1:500), and GAPDH (1:1000) primary antibodies (Santa Cruz Biotech) diluted in 5% BSA in PBS-T overnight at 4° C. The blots were then washed 2 × with PBS-T for 5 min each, and 1 × with PBS for 5 min, followed by incubation with goat anti-mouse secondary antibody (1:5000) diluted in 5% BSA in PBS-T for 1 h at room temperature. Finally, the blots were washed as before and incubated with Supersignal west pico chemiluminescence substrate (Pierce) and developed using ChemiDoc XRS+ (BioRad).

### Statistical analysis

Data are presented as the means of at least three independent experiments, and the error bars represent the standard deviation from the means. Statistical significance was calculated using one-way ANOVA. For statistical significance, *p-*value of <0.05 was used.

## Results and Discussion

### Endothelial cell vimentin expression and knock-down

The expression of vimentin in endothelial cells is shown in Figure 2. Endothelial cells were treated with different growth factors bFGF, EGF and VEGF at a 3 ng/mL concentration. Inhibition of vimentin filament formation is accomplished by adding 5 µM FiVe1 or transfecting endothelial cells with vimentin siRNA. The data showed that the ECs cultured in the presence of bFGF grow in close contact with each other similar to the control. On the other hand, cells treated with EGF or VEGF were separated, which may indicate endothelial-to-mesenchymal (EndMT) transition due to loss of cell-cell contact^19^. This transition is important for the formation of new microvasculature^20^. Furthermore, ECs treated with VEGF and EGF exhibit an elongated shape of the vimentin filament network which might also be an indication of the EndMT transition^19^. The addition of Five1 into the culture or transfection of ECs with vimentin siRNA disrupts the vimentin fibers assembly. As expected, the level of expression of vimentin is also reduced in the transfected ECs.

**Figure 2.**
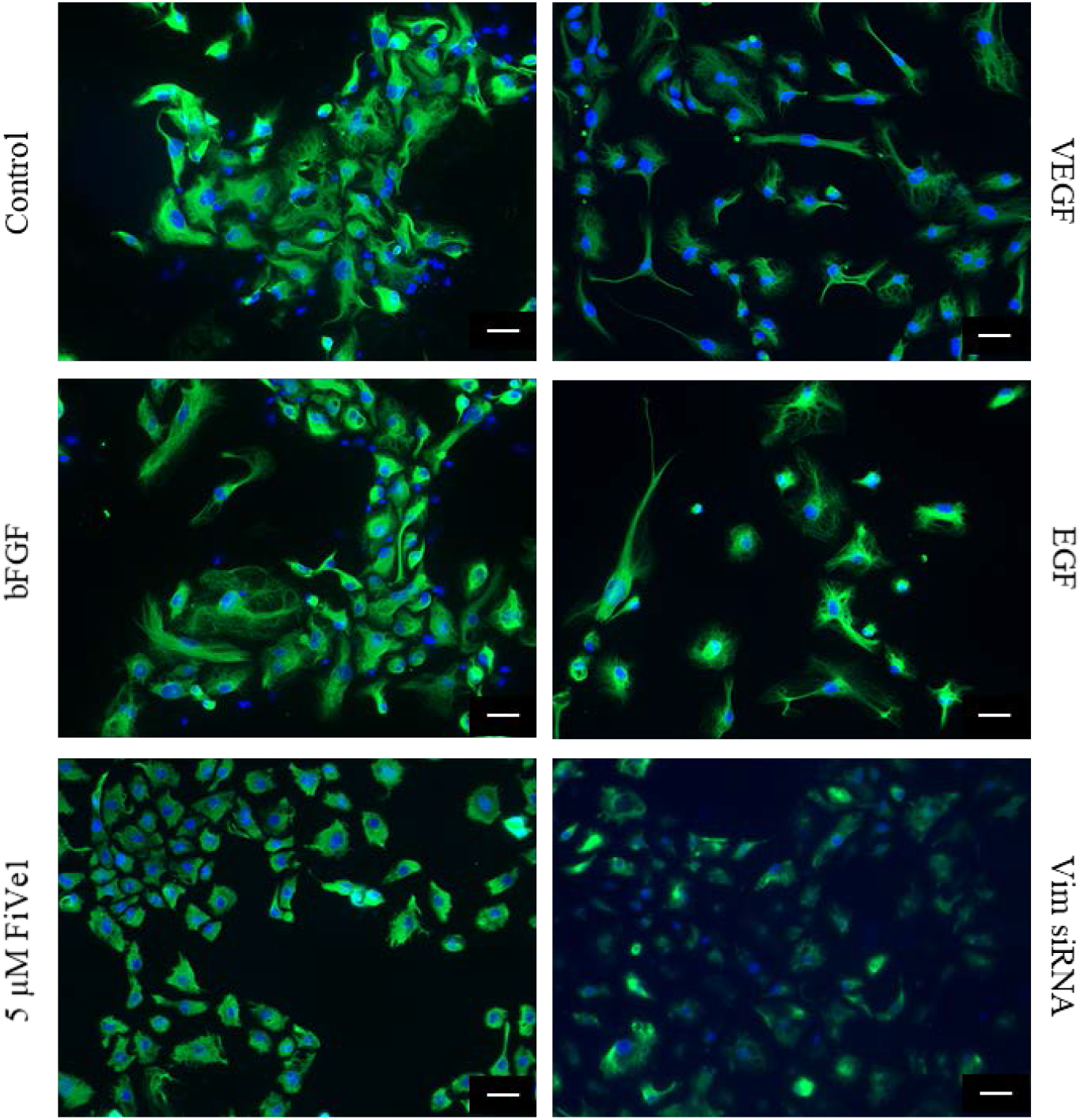
Fluorescence microscopy of expression of vimentin in HCAEC. HCAEC were treated with 3ng/mL of each bFGF, EGF, and VEGF. Vimentin filament disruption is shown by the addition of 5 µM FiVe1 or vimentin knock-down using vimentin siRNA. Scale bar = 50 µm.

### Role of vimentin in Notch signaling in static and dynamic cultures

To assess the role of vimentin in the regulation of the Notch signaling pathway, 10T1/2 cells were co-cultured with endothelial cells (EC), or ECs treated with VEGF, or ECs transfected with the vimentin siRNA. Figure 3. A shows the expression of three Notch target genes, Hes-1; Hes-5 and Acta2, under static conditions. Hes-1 was upregulated in the presence of ECs regardless of EC treatment conditions. Hes-5 expression was also upregulated in the presence of ECs; however, its expression was reduced slightly when vimentin expression was knocked down with siRNA; but it was still higher than control levels. The expression of Acta2 increased in the presence of ECs and EC (siRNA) and it was further increased in the VEGF treated ECs. A similar trend is observed for the protein expression of Hes-5 and Acta in figure 5.3.C. These results show that in static conditions, only the expression of Hes-5 was affected by the knock-down of vimentin expression.

**Figure 3.**
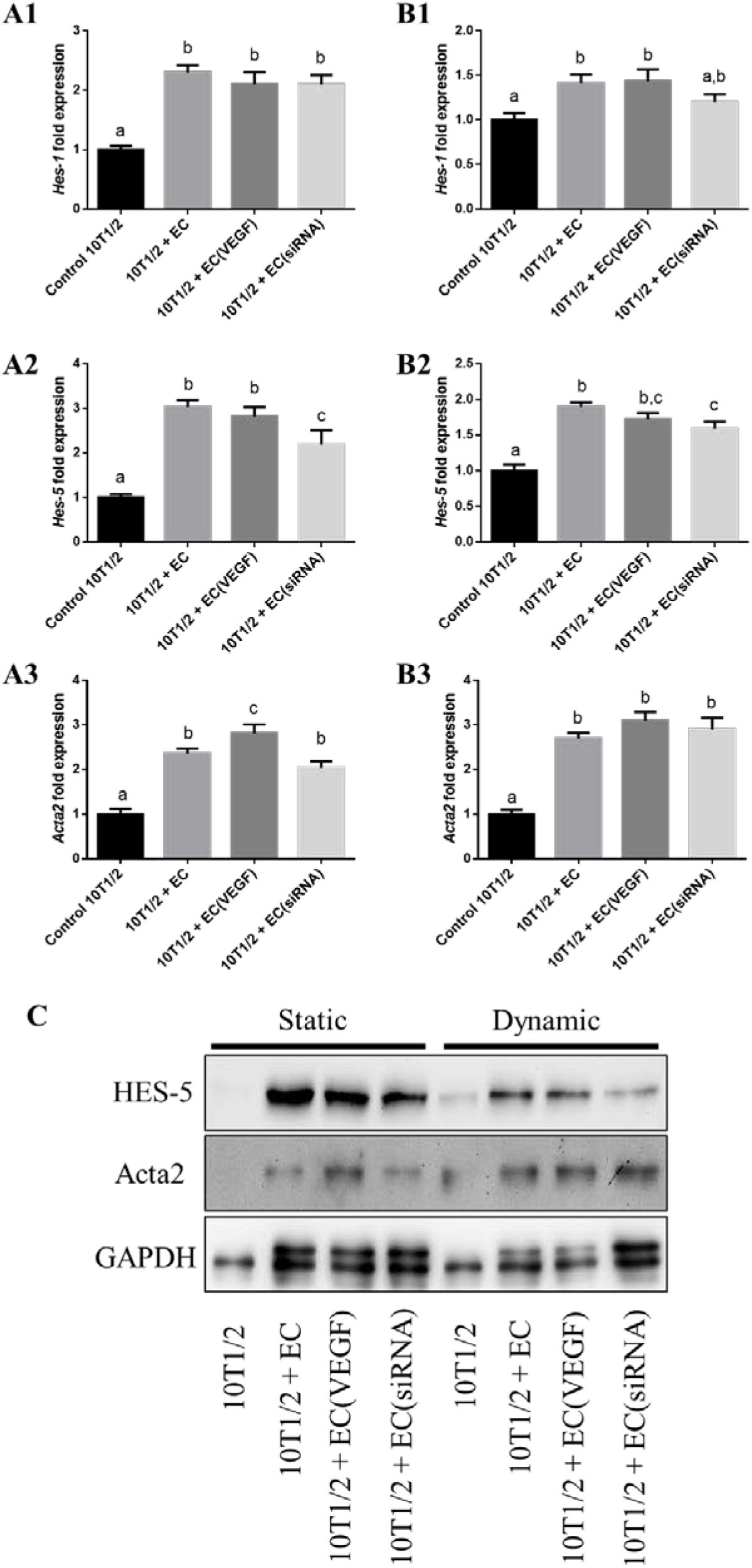
Co-culture of 10T1/2 and HCAEC in static and dynamic conditions. **(A)** Hes-1, Hes-5 and Acta2 gene expression of 10T1/2 cells co-cultured with EC, EC treated with VEGF (EC(VEGF)) or EC transfected with vimentin siRNA (EC (siRNA)) in static culture conditions. **(B)** Hes-1, Hes-5, and Acta2 gene expression of 10T1/2 cells co-cultured with EC, EC(VEGF), and EC (siRNA) in dynamic culture conditions. Lowercase letters on top of the bars are used for statistical analysis. Different lowercase letters indicate a statistically significant difference (p<0.05), whereas the same letters indicate no statistical difference (p>0.05). **(C)** Western blot analysis of Acta2 and Hes5 proteins in 10T1/2 cells co-cultured with EC, EC(VEGF), or EC (siRNA); GAPDH was used as a loading control.

Due to its tight interaction with integrins, vimentin functions as a mechanosensor of external forces from outside the cell and relays them to the nucleus allowing the cells to respond to such forces^21^. Due to this, the static culture conditions were repeated in dynamic conditions to assess whether shear forces might have an effect on the role of vimentin in regulating the Notch signaling. Under dynamic conditions, the expression of the Hes-1 gene increased in the presence of EC and was slightly reduced when vimentin was knocked-down by siRNA (Figure 3B1). A similar pattern is observed for the Hes-5 gene in Figure 3B2. On the other hand, the expression of Acta2 is upregulated in the presence of EC regardless of EC treatment (Figure 3.B3). These patterns are reflected at the protein level shown by the Western blot (Figure 3C.). These results show that vimentin might play a partial role in the Notch signal regulation because there was a slight decrease in the expression of Hes-5 when vimentin was knocked down, even though the other genes were not downregulated. Interestingly, Figure 3C shows that while the expression of Hes-5 was generally lower in dynamic conditions, a comparison between the control groups of static vs. dynamic cultures shows a slight upregulation of the Hes-5 gene. On the other hand, the expression of Acta2 appears to be generally upregulated in dynamic conditions. This might suggest that shear force alone plays a role in regulating Notch even though it might not be through vimentin. The cytoskeleton is a complex network of interconnected microfilaments, intermediate filaments, and microtubules and therefore, there might be a redundancy in the role of the cytoskeleton in regulating the Notch signal. Thus, despite the fact that the vimentin expression was knocked down, other cytoskeletal components might compensate to keep vital signaling pathways operational.

### Vimentin filament disruption partially affects Notch signaling

The effect of vimentin filament disruption was tested on the Hes-1, Hes-5, and Acta2 gene expression levels in 10T1/2 cells co-cultured with HCAECs in 2D cultures. Figure 4A shows that disruption of vimentin filament networks did not significantly affect the gene expression of Hes-1 or Hes-5 Notch target genes even though there was a downward trend in the expression of both these genes in the FiVe1 treated group. On the other hand, while Five1 treatment reduced the gene expression of the Acta2 gene, its expression was still higher than the control. Conversely, Acta2 protein expression levels remained high in 3D co-culture even in the presence of FiVe1-treated HCAECs, as shown in Figure 4B. This data shows that while vimentin filament disruption slightly decreased the gene expression of Notch downstream targets, its effect is minimal and does not have a noticeable effect on the protein expression level (Figure 4B). This could be due to the redundant role of intermediate filaments such as vimentin, where the disruption of one filament network is compensated by other elements in the cytoskeleton.

**Figure 4.**
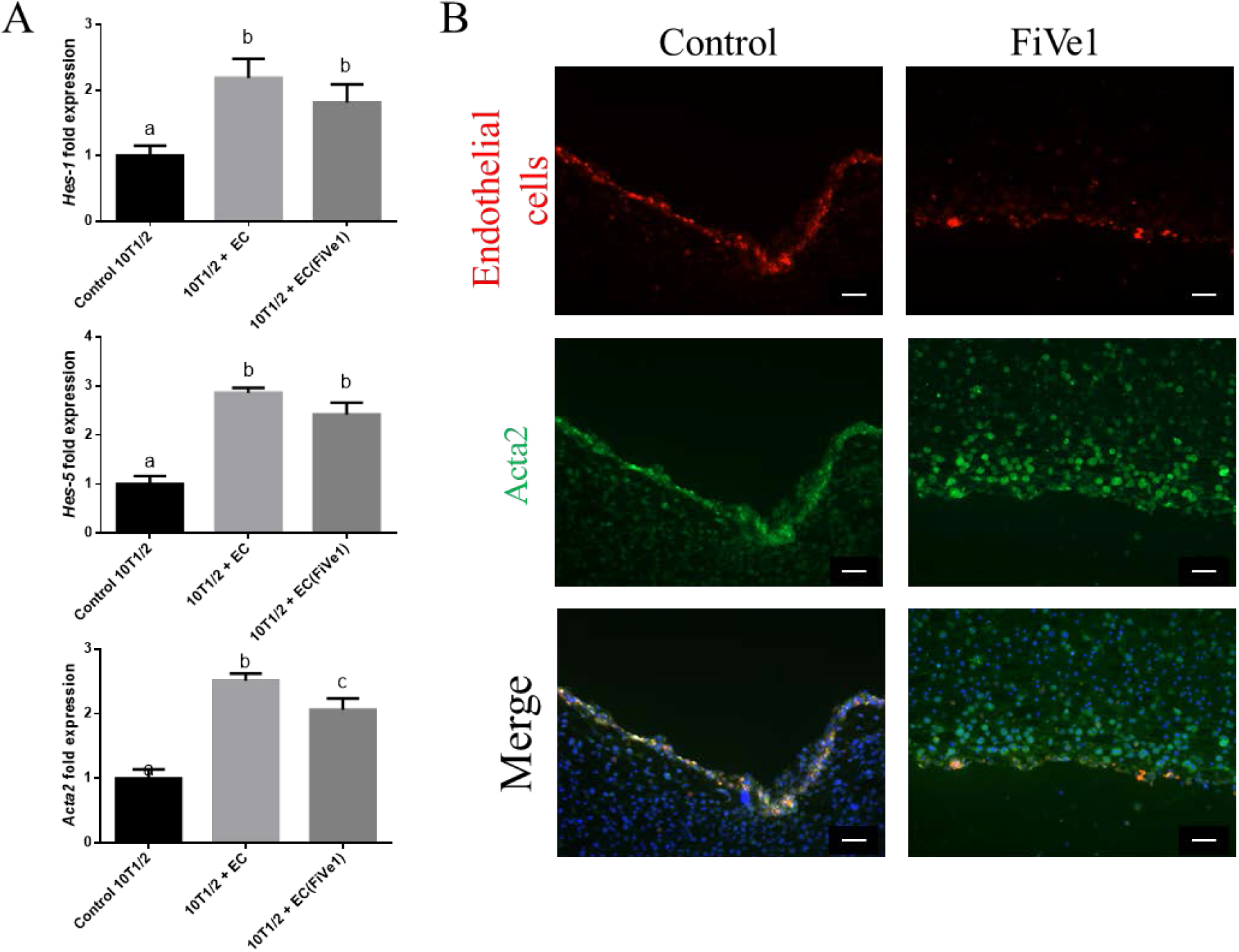
Gene and protein expression of 10T1/2 co-cultured with HCAEC in 2D and 3D, respectively. **(A)** Hes-1, Hes-5 and Acta2 gene expression in 10T1/2 cells co-cultured with HCAECs on 2D. **(B)** Immunofluorescence microscopy showing expression of Acta2 in 3D co-culture of 10T1/2 and HCAEC in control *vs*. FiVe1 treated tissues. Scale bar = 50 µm.

**Figure 5.**
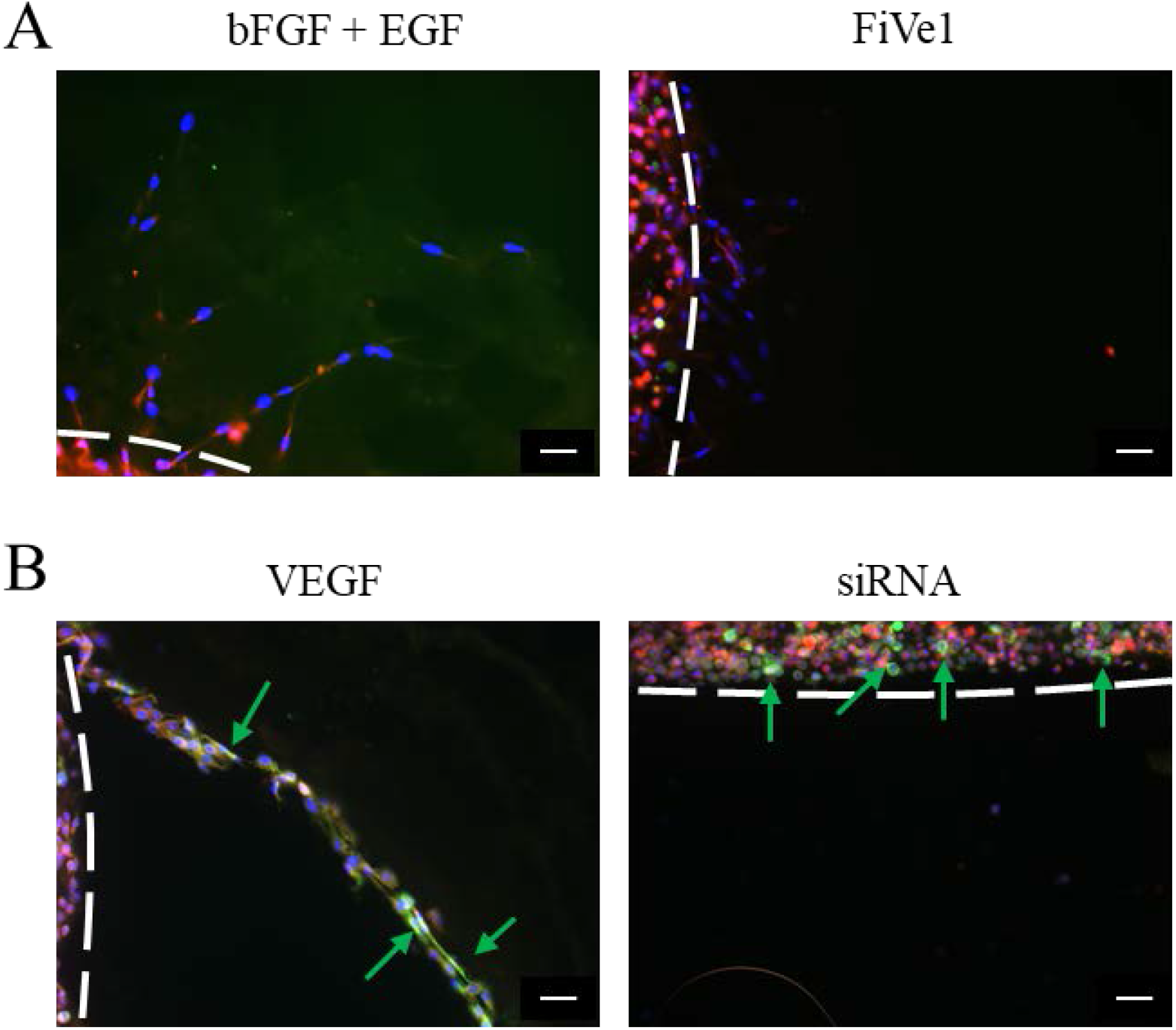
Endothelial cell migration and neovascularization in 3D fibrin gel. (A) Migration of endothelial cells into low-concentration fibrin gel in the presence of bFGF and EGF and FiVe1 treated gels. (B) Micro-vessel formation in 3D fibrin gels containing VEGF, or HCAECs that were previously transfected with vimentin siRNA. Scale bar = 50 µm. Dashed white lines indicate tissue outline. The green arrows indicate Acta2 positive cell**s**.

### Vimentin plays a role in endothelial cell migration and micro-vessel formation

The ability of endothelial cells to migrate in a 3D environment was tested in response to angiogenic growth factors (bFGF + EGF or VEGF). Furthermore, the effect of vimentin disruption on this process was tested using a small molecule inhibitor, FiVe1, or gene expression knock-down using vimentin siRNA (Figure 5.). The results showed that angiogenic growth factors were able to induce the migration of endothelial cells out of the 3 mg/mL fibrin gel and into the less concentrated 1 mg/mL fibrin gel, which contained the growth factors (Figure 5). On the other hand, when vimentin filament fiber formation was inhibited, migration was greatly affected. Furthermore, while bFGF and EGF combination has been shown to induce growth and proliferation in endothelial cells^22^, VEGF is a notably more potent angiogenic factor since the sprouting micro-vessels are better organized (Figure 5B). Moreover, these micro-vessels were able to recruit 10T1/2 cells that expressed Acta2 as shown by the green staining indicated by the arrows. In contrast, siRNA-transfected HCAECs were not able to migrate out of the 3 mg/mL fibrin; however, they were able to induce expression of Acta2 in 10T1/2 cells, which is in agreement with the cells treated with VEGF. Taken together, this exploratory study results showed that vimentin plays an important role in the migration of endothelial cells and recruits 10T1/2 cells to form micro-vessels.

## Conclusion

In this exploratory study, the role of vimentin in the regulation of Notch signaling and neovascularization was explored. The results demonstrated that vimentin plays a limited role in Notch signaling, as evidenced by a slight reduction of certain Notch signaling targets (Hes-5 and Acta2) when vimentin filaments were inhibited. However, vimentin plays a much more important role in micro-vessel formation, which allows endothelial cells to migrate and recruit 10T1/2 cells in response to VEGF. While these pilot results shed some light on the role of vimentin in Notch signaling and neovascularization, more research is required to fully elucidate the role of this intermediate fiber in the context of vascular development.

